# Prenatal Bisphenol A Exposure in Mice Induces Multi-tissue Multi-omics Disruptions Linking to Cardiometabolic Disorders

**DOI:** 10.1101/336214

**Authors:** Le Shu, Qingying Meng, Brandon Tsai, Graciel Diamante, Yen-Wei Chen, Andrew Mikhail, Helen Luk, Beate Ritz, Patrick Allard, Xia Yang

**Affiliations:** Department of Integrative Biology and Physiology, University of California, Los Angeles, CA 90095, USA; Molecular, Cellular, and Integrative Physiology Interdepartmental Program, University of California, Los Angeles, CA 90095, USA; Molecular Toxicology Interdepartmental Program, University of California, Los Angeles, CA 90095, USA; Department of Epidemiology, Fielding School of Public Health, University of California, Los Angeles, CA 90095, USA; Institute for Society and Genetics, University of California, Los Angeles, CA 90095, USA; Institute for Quantitative and Computational Biosciences, University of California, Los Angeles, CA 90095, USA

**Author notes:** Corresponding author: (XY) Address: Department of Integrative Biology and Physiology, University of California, Los Angeles Terasaki Life Sciences Building 2000D Los Angeles, CA 90095. These authors contributed equally to this work (Le Shu, Qingying Meng).

## Abstract

The health impacts of endocrine disrupting chemicals (EDCs) remain debated and their tissue and molecular targets are poorly understood. Here, we leveraged systems biology approaches to assess the target tissues, molecular pathways, and gene regulatory networks associated with prenatal exposure to the model EDC Bisphenol A (BPA). Prenatal BPA exposure led to scores of transcriptomic and methylomic alterations in the adipose, hypothalamus, and liver tissues in mouse offspring, with cross-tissue perturbations in lipid metabolism as well as tissue-specific alterations in histone subunits, glucose metabolism and extracellular matrix. Network modeling prioritized main molecular targets of BPA, including *Pparg, Hnf4a, Esr1, Srebf1*, and *Fasn*. Lastly, integrative analyses identified the association of BPA molecular signatures with cardiometabolic phenotypes in mouse and human. Our multi-tissue, multi-omics investigation provides strong evidence that BPA perturbs diverse molecular networks in central and peripheral tissues, and offers insights into the molecular targets that link BPA to human cardiometabolic disorders.

## Introduction

A central concept in the Developmental Origins of Health and Disease (DOHaD) states that adverse environmental exposure during early developmental stages is an important determinant for later onset adverse health outcomes, even in the absence of continuous exposure in adulthood [1-3]. BPA is one of the most prevalent environmental metabolic disruptors identified to date with widespread exposure in human populations and likely plays a role in DOHaD. BPA is used in the production of synthetic polymers, including epoxy resins and polycarbonates [4]. The advantageous mechanical properties of BPA have resulted in its ubiquitous use in everyday goods such as plastic bottles and inner coating of canned foods [5, 6]. BPA exposure has been confirmed in the majority of human populations [7] and has been linked to body weight, obesity, insulin resistance, diabetes, metabolic syndrome (MetS), and cardiovascular diseases in both human epidemiologic and animal studies [8-15]. Importantly, it has been suggested that the developing fetus is particularly vulnerable to BPA exposure [8, 16]. Intrauterine growth retardation (IUGR) has been consistently observed after developmental BPA exposure at intake doses below the suggested human safety level and has been associated with low birth weight, elevated adult fat weight and altered glucose homeostasis [8, 17-20]. As a result, BPA has been banned from baby products in Europe, Canada, and the US. However, BPA is still in use in non-baby products, renewing concerns with regards to the continuous exposure of populations in addition to the description of its ability to influence health outcomes, including obesity and MetS, over several generations [21-24]. Together these lines of evidence support an intriguing hypothesis that BPA may have been a contributing factor to the rise of MetS and cardiometabolic diseases worldwide in the past decades [25-27].

Despite numerous studies connecting BPA with adverse health outcomes, there remain ample conflicting findings, as summarized by the European Food Safety Agency [28] and the BPA Joint Emerging Science Working Group of the US FDA. Although inconsistencies across studies might be attributable to non-monotonic dose response, exposure window difference, and varying susceptibility between testing models [13, 29], there are also several additional layers of complexity and challenges hindering the full dissection of the biological effects of BPA. First, previous studies examining BPA in various cell types and tissues suggest a broad impact on biological systems [23, 30-32]. Second, BPA has been found to modulate multidimensional molecular events, such as gene expression and epigenetic changes, that are functionally important for processes such as metabolism and immune response [33-38]. However, due to most studies being designed to focus on one factor at a time as well as non-comparable study designs, it is difficult to directly compare effects across tissues or types of molecular data to derive the molecular rules of sensitivity to BPA exposures. In a recent National Toxicology Program report, CLARITY-BPA, where multiple organs were examined, evidence of weight gain and cardiac dysfunctions were observed, however, the study was designed to be solely descriptive and no mechanism of action was proposed. These research gaps in our understanding of the pleiotropy of EDCs and toxicant biological actions necessitated the establishment of the NIEHS TaRGET consortium and a more recent call for the research community to systemically interrogate multiple omics in multiple tissues to accelerate the discovery of key biological fingerprints of environmental exposure [39].

Here, we address some of the aforementioned limitations of past studies by using a highly integrative approach. We conducted a multi-tissue, multi-omics systems biology study to examine the systems level influence of prenatal BPA exposure using modern integrative genomics and network modeling approaches in a mouse model.

We first utilized next-generation sequencing technologies to characterize perturbations in both the transcriptome and the epigenome across three tissues (white adipose tissue, hypothalamus, liver) in mouse offspring who had experienced *in utero* exposure to BPA. Based on mounting evidence that genes operate in highly complex tissue-specific regulatory networks, we hypothesized that prenatal BPA exposure induces genomic and epigenomic reprogramming in the offspring by affecting the organization and function of tissue-specific gene networks [40-43]. Using both transcription factor (TF) networks and Bayesian networks, we modeled the dynamics of transcriptomic and epigenomic signatures and predicted potential regulators that govern the actions of BPA. Furthermore, the transcriptome, epigenome, and network information were layered upon metabolic phenotypes such as body weight, adiposity, circulating lipids, and glucose levels in the mouse offspring to evaluate disease association. Lastly, to assess the relevance of the BPA molecular targets identified in our mouse model for human diseases, we applied integrative genomics to bridge the mouse molecular signatures and genetic disease association data from human studies. Our study represents a comprehensive data-driven, systems-level investigation of the molecular and health impact of BPA.

## Results

### Prenatal BPA exposure induces intrauterine growth retardation (IUGR) and alterations in cardiometabolic phenotypes

As shown in **Fig 1A**, pregnant C57BL/6 mice were exposed to BPA during gestation (day 1 to day 20 post-conception) via oral gavage at the dosage of 5mg/kg/day, situated below most reported no-observed-adverse-effect-level (NOAEL) according to toxicity testing (https://comptox.epa.gov/dashboard/dsstoxdb/results?search=Bisphenol+A). This dosage was typically used in previous studies [23, 44-46], and was chosen as a proof-of-concept for our systems biology study design and to facilitate comparison with previous studies. Male and female offspring (n = 9 for control and n = 11 for BPA in male; n = 9 for control and n = 13 for BPA in female, 2-3 mice from 3-4 litters/group; **S1 Fig**) of weaning age (3-weeks) were examined for a spectrum of metabolic phenotypes (detailed below). We chose the weaning age in order to investigate early molecular and phenotypic changes in the offspring, which may predispose the offspring to late onset diseases. Compared with the control group, both male and female offspring from the BPA group showed significantly lower body weight, indicative of IUGR, a trait that is strongly associated with later life insulin resistance and obesity risk (**Fig 1B, D**). There were also significant decreases in serum lipid parameters and an increase in serum glucose level in males (**Fig 1C**), but not in females (**Fig 1E**). The decreases in the lipid parameters at this early developmental stage likely reflect the growth retardation phenotype observed and may provide feedback signals to predispose the exposed offspring to lipid dysregulation later in life. The phenotypic differences between BPA and control groups are not the results of litter effect, as offspring from different dams in each group showed similar patterns (**S1 Fig**).

**Fig 1.**
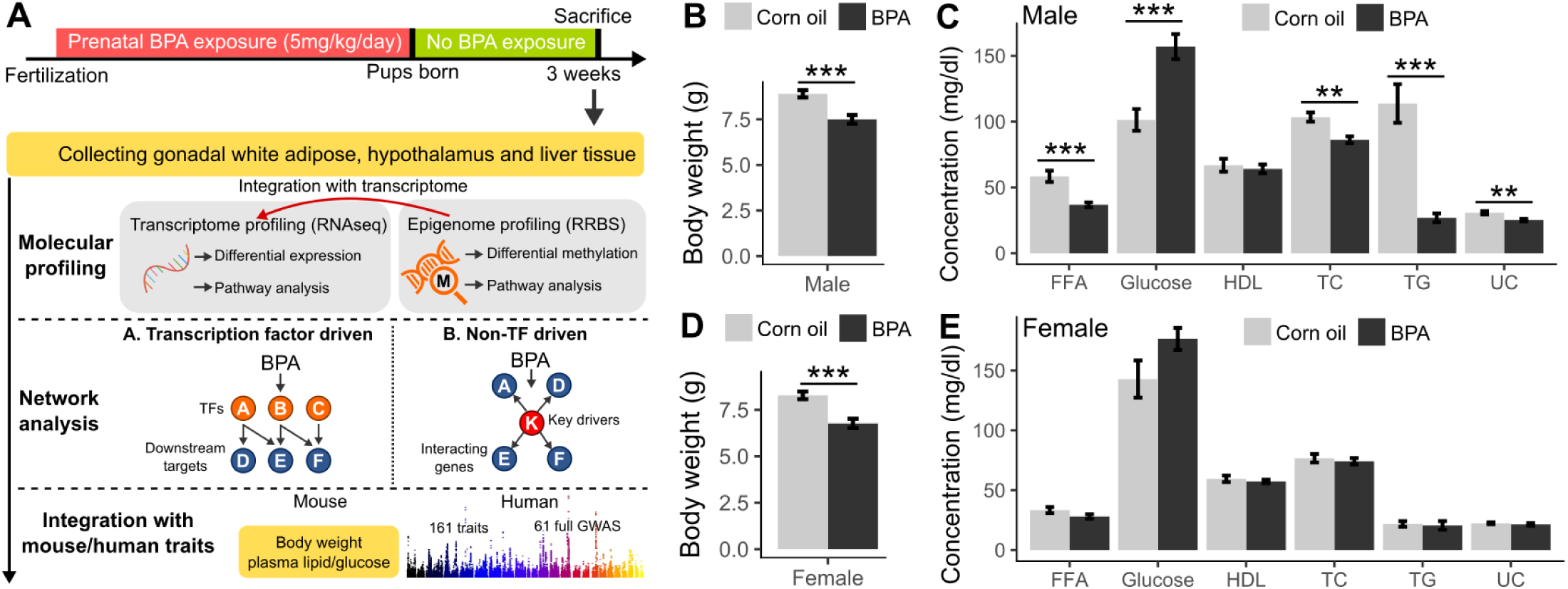
Overall study design and the measurements of metabolic traits in male and female offspring. (A) Framework of multi-omics approaches to investigate the impact of prenatal BPA exposure. B-C) Comparison of body weight, serum lipids and glucose level in male mice at weaning age. D-E) Comparison of body weight, serum lipids and glucose level in female mice at weaning age. FFA: free fatty acid; HDL: high-density lipoprotein cholesterol; TC: total cholesterol; TG: triglyceride; UC: unesterified cholesterol. * p < 0.05, ** p < 0.01, *** p < 0.001 by two-sided Student’s T-test. N=9-13 mice (3-4 litters from different dams)/group.

### Prenatal BPA exposure induces tissue-specific transcriptomic alterations in male weaning offspring

To explore the molecular basis underlying the potential health impact of prenatal BPA exposure, we collected three key metabolic tissues including white adipose tissue, hypothalamus, and liver from male offspring (due to the stronger observed phenotypes) at 3 weeks. The hypothalamus is the central regulator of endocrine and metabolic systems, whereas liver and white adipose tissues are critical for energy and metabolic homeostasis. We used RNA sequencing (RNA-seq) to profile the transcriptome, and identified 86, 93, and 855 differentially expressed genes (DEGs) in the adipose tissue, hypothalamus, and liver tissue respectively, at false discovery rate (FDR) < 0.05 (**Fig 2A, S1 Table**). This supports the ability of prenatal BPA exposure to induce large-scale transcriptomic disruptions in offspring, with the impact appearing to be more prominent in liver. The DEGs were highly tissue-specific, with only 12 out of the 86 adipose DEGs and 16 out of the 93 hypothalamus DEGs being found in liver. Interestingly, the hypothalamic DEGs are predominantly up-regulated in the BPA group whereas the other two tissues did not show such direction bias (**S2 Table**). Only one gene, *Cyp51* (sterol 14-alpha demethylase), was shared across all three tissues but with different directional changes (upregulated in hypothalamus and liver, downregulated in adipose) (**Fig 2B**). The Cyp51 protein catalyzes metabolic reactions including cholesterol and steroid biosynthesis and biological oxidation [47] and is a critical regulator for testicular spermatogenesis [48]. The consistent alteration of *Cyp51* across tissues suggests that this gene is a general target of BPA, with the potential to alter functions related to cholesterol, hormone, and energy metabolism.

**Fig 2.**
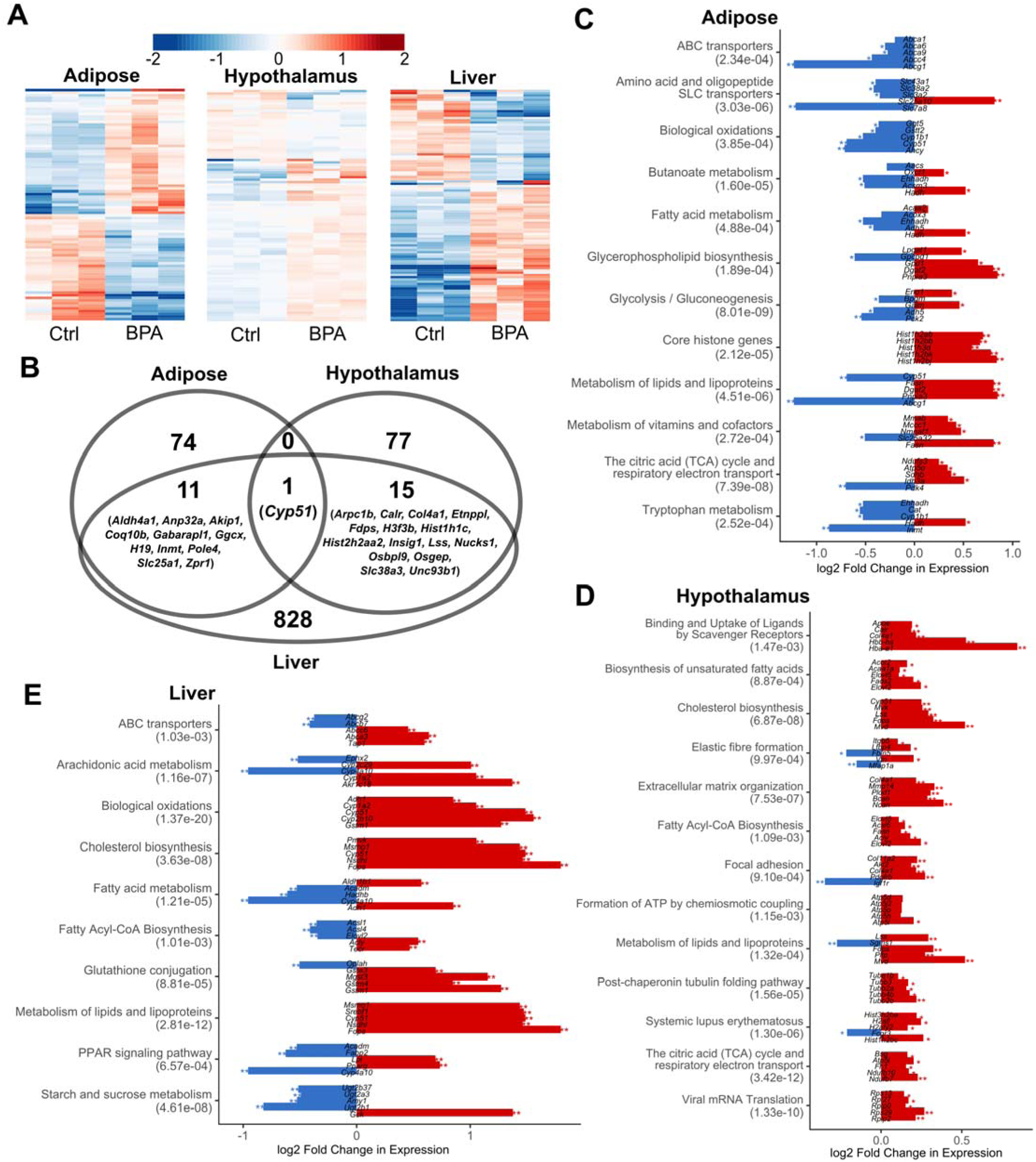
Prenatal BPA exposure induced transcriptomic alterations in adipose, hypothalamus and liver. (A) Heatmap of expression changes in adipose, hypothalamus and liver for the top 100 differentially expressed genes (DEGs) affected by BPA. Color indicates fold change of expression, with red and blue indicating upregulation and downregulation by BPA. (B) Venn Diagram demonstrating tissue-specific and shared DEGs between tissues. (C-E) Significantly enriched pathways (FDR < 5%) among DEGs from each tissue. Enrichment p-value (shown in parenthesis following the name of functional annotation) is determined by MSEA. The fold change and statistical significance for the top 5 differentially expressed genes in each pathway are shown. *, p < 0.05; **, FDR < 5% in differential expression analysis using DEseq2.

### Functional annotation of DEGs in adipose, hypothalamus, and liver tissues

To better understand the biological implications of the BPA exposure related DEGs in individual tissues, we evaluated the enrichment of DEGs for known biological pathways and functional categories using the Mergeomics package [49] (**Fig 2C-E,** full results in **S3 Table**). We observed strong enrichment for pathways related to lipid metabolism (lipid transport, fatty acid metabolism, cholesterol biosynthesis) and energy metabolism (biological oxidation, TCA cycle) across all three tissues. Most of these pathways appeared to be upregulated in all three tissues, with the exception of downregulation of genes involved in biological oxidation in adipose tissue (**Fig 2C-E**). Individual tissues also showed perturbations of unique pathways: PPAR signaling and arachidonic acid pathways were altered in liver; extracellular matrix related processes were enriched among hypothalamic DEGs; core histone genes were upregulated in adipose DEGs (**Fig 2C-E**). In addition, triglyceride biosynthesis and glucose metabolism pathways were also moderately enriched among adipose DEGs, whereas few changes were seen for genes involved in adipocyte differentiation (**S2 Fig**).

### Replication of the DEG signatures using independent studies

To support the replicability of the differential expression signatures identified in our study, we sought to validate the signatures using independent expression profiling data deposited on GEO (**S1 Text**, Supplemental Methods). We identified three GEO datasets - two from GSE26728 [50] and one from GSE43977 [51] (**S3A Fig**) - that characterized the liver transcriptome following BPA exposure during adulthood. Highlighting the novelty of our study, we were not able to identify other datasets with the same *in utero* exposure condition tested in our exposure paradigm, making a direct replication difficult. However, we reasoned that if core mechanisms exist for BPA regardless of experimental conditions, consistent signals should be derived. We compared the differential expression signatures from the three existing liver studies against ours, and found limited consistency in BPA signatures across datasets, even for the two datasets that were originated from the same study (GSE26728) (**S3B Fig**). These results support that BPA has condition-specific activities. Nevertheless, 10% of our DEGs were replicated in the other GEO datasets (p < 1e-4 compared to random expectation via a permutation analysis, **S3C Fig**). *Srebf1* (Sterol Regulatory Element Binding Transcription Factor 1), a key transcription factor in lipid metabolism, was consistent across all four datasets, along with numerous additional genes consistent in two or more studies (**S3C Fig**).

Next, we compared the enriched pathways among the DEGs from each dataset to evaluate whether distinct study-specific signatures could converge onto similar biological processes. The replicated pathways across studies include steroid hormone biosynthesis, retinol metabolism and fatty acid metabolism, suggesting that these processes were consistently influenced by BPA under varying exposure windows and dosages (**S3D Fig**). At FDR < 5%, 56.1% of the significant pathways in our study were replicated in one or more independent studies (p < 1e-4 compared to random expectation via a permutation test, **S3E Fig**). Pair-wise comparison revealed relatively higher overlap ratios between our study and individua independent studies than between the previous studies, despite the greater similarity in the study design among the previous studies (**S3E Fig**). Overall, the generally higher pair-wise replication rates of the biological pathways between our study and independent studies support the adequate power of our study to reveal core BPA-associated biological pathways implicated in previous studies.

### Prenatal BPA exposure induces tissue-specific epigenetic alterations in male weaning offspring

Consistent with the observed gene expression disruptions at the transcriptomic level, we observed numerous methylomic alterations using reduced representation bisulfite sequencing (RRBS), which characterizes DNA methylation states of millions of potential epigenetic sites at single base resolution. At FDR < 5%, 5136, 104, and 476 differentially methylated CpGs (DMCs) were found in adipose, hypothalamus, and liver tissues, respectively (**Fig 3A, S4 Table**). When comparing our adipose methylation signatures with a previous study [36], we were able to replicate 5 out of 7 peak hypomethylated genes, and 6 out of 9 peak hypermethylated genes. Interestingly, BPA induced local methylation changes in *Gm26917* and *Yam1*, two long non-coding RNAs (lncRNAs) with no previously known link to BPA, consistently across three tissues (**Fig 3B**). The majority of the DMCs are located in intergenic regions (32% - 38%), followed by introns (31% - 37%) and exons (13% - 15%), but there is a paucity of DMCs in the promoter region (3% - 5%) (**S4 Fig**). Contrary to predictions that promoter regions may be more prone to epigenetic changes, we found that within-gene and intergenic methylation alterations in DNA methylation are more prevalent, a pattern consistently observed in previous epigenomic studies [41, 52]. In addition, 5.0%, 8.6%, and 8.1% DMCs overlap with repetitive DNA elements in adipose, hypothalamus, and liver, respectively, recapitulating previous report of the interaction between BPA and repetitive DNA [53].

**Fig 3.**
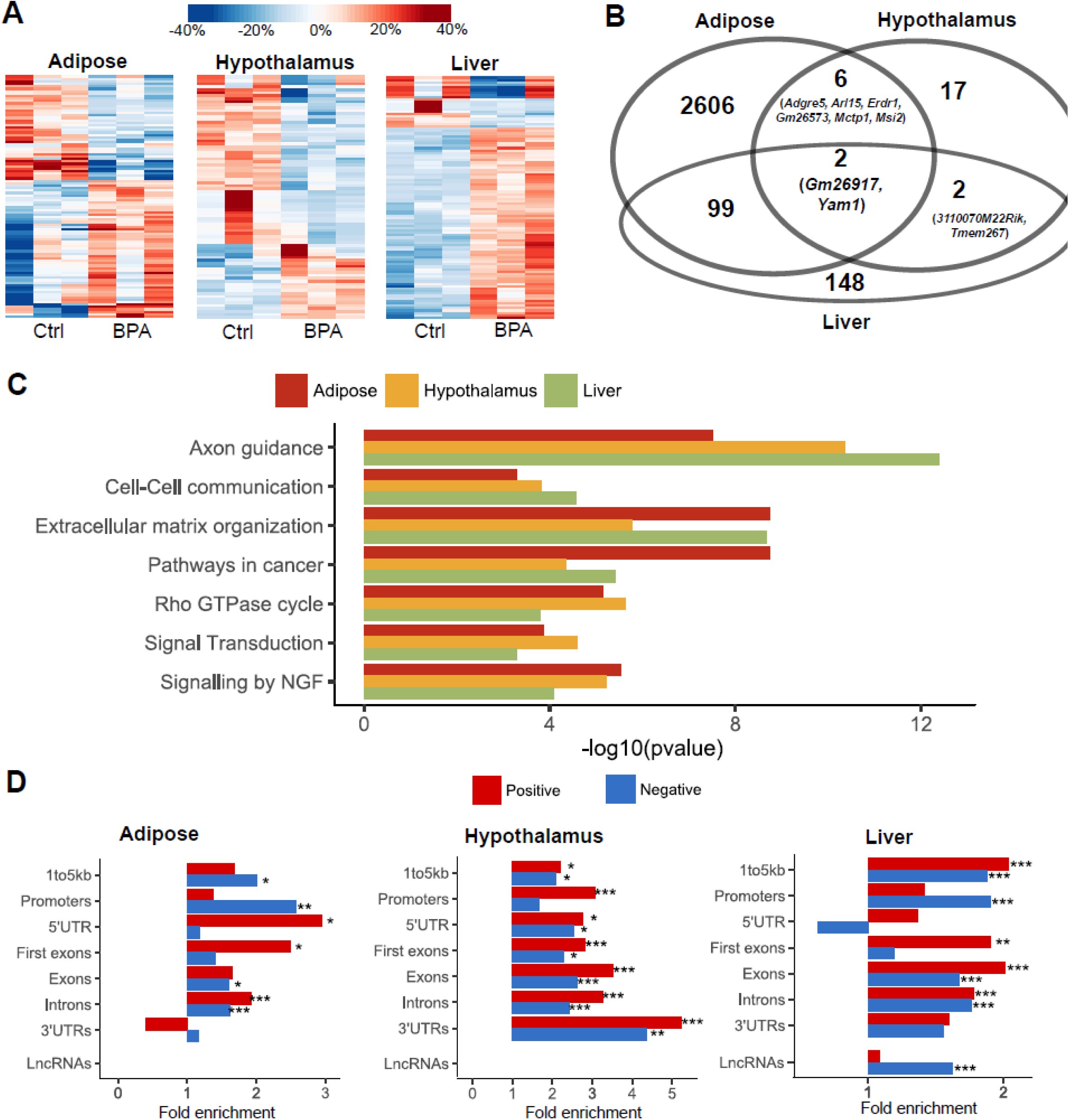
Prenatal BPA exposure induced methylomic level alteration in adipose, hypothalamus and liver. (A) Heatmap of methylation level changes for the top 100 differentially methylated CpGs (DMCs). Color indicates change in methylation ratio, with red and blue indicating upregulation and downregulation by BPA. (B) Venn Diagram of genes with local DMCs between tissues shows tissue-specific and shared genes mapped to DMCs. (C) Significantly enriched pathways that satisfied FDR < 1% across DMCs from adipose, hypothalamus, and liver tissues. Enrichment p-value is determined by MSEA. (D) Fold enrichment for positive correlations (red bars) or negatively correlations (blue bars) between DMCs and local DEGs, assessed by different gene regions. *, p < 0.05; **, p < 0.01; ***, p < 0.0001; enrichment p-values were determined using Fisher’s exact test.

For DMCs that are located within or adjacent to genes, we further tested whether the local genes adjacent to those DMCs show enrichment for known functional categories. Unlike DEGs, top processes enriched for DMCs concentrated on intra- and extra-cellular communication and signaling related pathways such as axon guidance, extracellular matrix organization and NGF signaling (**Fig 3C**, **full results in S5 Table**). The affected genes in these processes are related to cellular structure, cell adhesion, and cell migration, indicating that these functions may be particularly vulnerable to BPA induced epigenetic modulation.

### Potential regulatory role of DMCs in transcriptional regulation of BPA induced DEGs

To explore the role of DMCs in regulating DEGs, we evaluated the connection between transcriptome and methylome by correlating the expression level of DEGs with the methylation ratio of their local DMCs. For the DEGs in adipose, hypothalamus and liver tissue, we identified 42, 36, and 278 local DMCs whose methylation ratios were significantly correlated with the gene expression. At a global level, compared to non-DEGs, DEGs are more likely to contain local correlated DMCs (**S5 Fig**). A closer look into the expression-methylation correlation by different chromosomal regions further revealed a context dependent correlation pattern (**Fig 3D**). In adipose and liver, the 3-5% of DMCs in promoter regions tend to show significant enrichment for negative correlation with DEGs, whereas gene body methylations for DEGs are more likely to show significant enrichment for positive correlation with gene expression. In hypothalamus, however, positive correlations between DEGs and DMCs are more prevalent across different gene regions. In addition, liver DMCs within lncRNAs were uniquely enriched for negative correlation with lncRNA expression, although the lack of a reliable mouse lncRNA target database prevented us from further investigating whether downstream targets of the lncRNAs were enriched in the DEGs. Specific examples of DEGs showing significant correlation with local DMCs include adipose DEG *Slc25a1* (Solute Carrier Family 25 Member 1, involved in triglyceride biosynthesis), hypothalamic DEG *Mvk* (Mevalonate Kinase, involved in cholesterol biosynthesis), and liver DEG *Gm20319* (a lncRNA with unknown function) (**S6 Fig and S6 Table**). These results support a role of BPA-induced differential methylation in altering the expression levels of adjacent genes.

### Pervasive influence of prenatal BPA exposure on the liver transcription factor network

BPA is known to bind to diverse types of nuclear receptors such as estrogen receptors and peroxisome proliferator-activated (PPAR) receptors that function as transcription factors (TFs), thus influencing the action of downstream genes [54, 55]. *PPARg* in particular has been shown to be a target of BPA in mouse and human and mechanistically linking BPA exposure with its associated effect on weight gain and increased adipogenesis [56-58]. To explore the TF regulatory landscape underlying BPA exposure based on our genome-wide data, we leveraged tissue-specific TF regulatory networks from the FANTOM5 project [59] and integrated it with our BPA transcriptome profiling data. No TF was found to be differentially expressed in adipose tissue, whereas 1 TF (Pou3f1) and 14 TFs (such as Esrra, Hnf1a, Pparg, Tcf21, Srebf1) were found to be differentially expressed in hypothalamus and liver, respectively. Due to the temporal nature of TF action, changes in TF levels may precede the downstream target genes and not be reflected in the transcriptomic profiles measured at the time of sacrifice. Therefore, we further curated the target genes of TFs from FANTOM5 networks and tested the enrichment for the target genes of each TF among our tissue-specific DEGs (**S7 Table**). This analysis confirmed that BPA perturbs the activity of the downstream targets for estrogen receptors Esrrg (p = 1.4e-3, FDR = 1.9%) and Esrra (p = 0.03, FDR = 13%) in liver, as well as Esr1 in both adipose (p = 7.2e-3, FDR = 10.6%) and liver (p = 7.2e-3, FDR = 4.7%). Targets of Pparg were also perturbed in liver (p = 4.1e-3, FDR = 3.8%). Therefore, we demonstrated that our data-driven network modeling is able to not only recapitulate results from previous *in-vitro* and *in-vivo* studies showing that BPA influences estrogen signaling and PPAR signaling [55], but also uniquely point to the tissue specificity of these BPA target TFs.

In addition to these expected TFs, we identified 14 adipose TFs and 61 liver TFs whose target genes were significantly enriched for BPA DEGs at FDR < 5%. Many of these TFs showed much stronger enrichment for BPA DEGs among their downstream targets than the estrogen receptors (**S7 Table**). The adipose TFs include nuclear transcription factor Y subunit alpha (Nfya) and fatty acid synthase (Fasn), both implicated in the adipocyte energy metabolism [60]. The liver TFs include multiple genes from the hepatocyte nuclear factors (HNF) family and the CCAAT-enhancer-binding proteins (CEBP) family, which are critical for liver development and function, suggesting a pervasive influence of BPA on liver TF regulation.

We further extracted the subnetwork containing 89 unique downstream targets of the significant liver TFs that are also liver DEGs. This subnetwork showed significant enrichment for genes involved in metabolic pathways such as steroid hormone biosynthesis and fatty acid metabolism. The regulatory subnetwork for the top liver TFs (FDR < 5%) revealed a highly interconnected TF subnetwork that potentially senses BPA exposure and in turn governs the expression levels of their targets (**Fig 4A**), with *Pparg* and *Hnf4* among the core TFs. Some of the TFs in this network, including *Esr1, Esrrg, Foxp1,* and *Tcf7l1*, also had local DMCs identified in our study, indicating that BPA may perturb this liver TF subnetwork via local modification of DNA methylation of key TFs.

**Fig 4.**
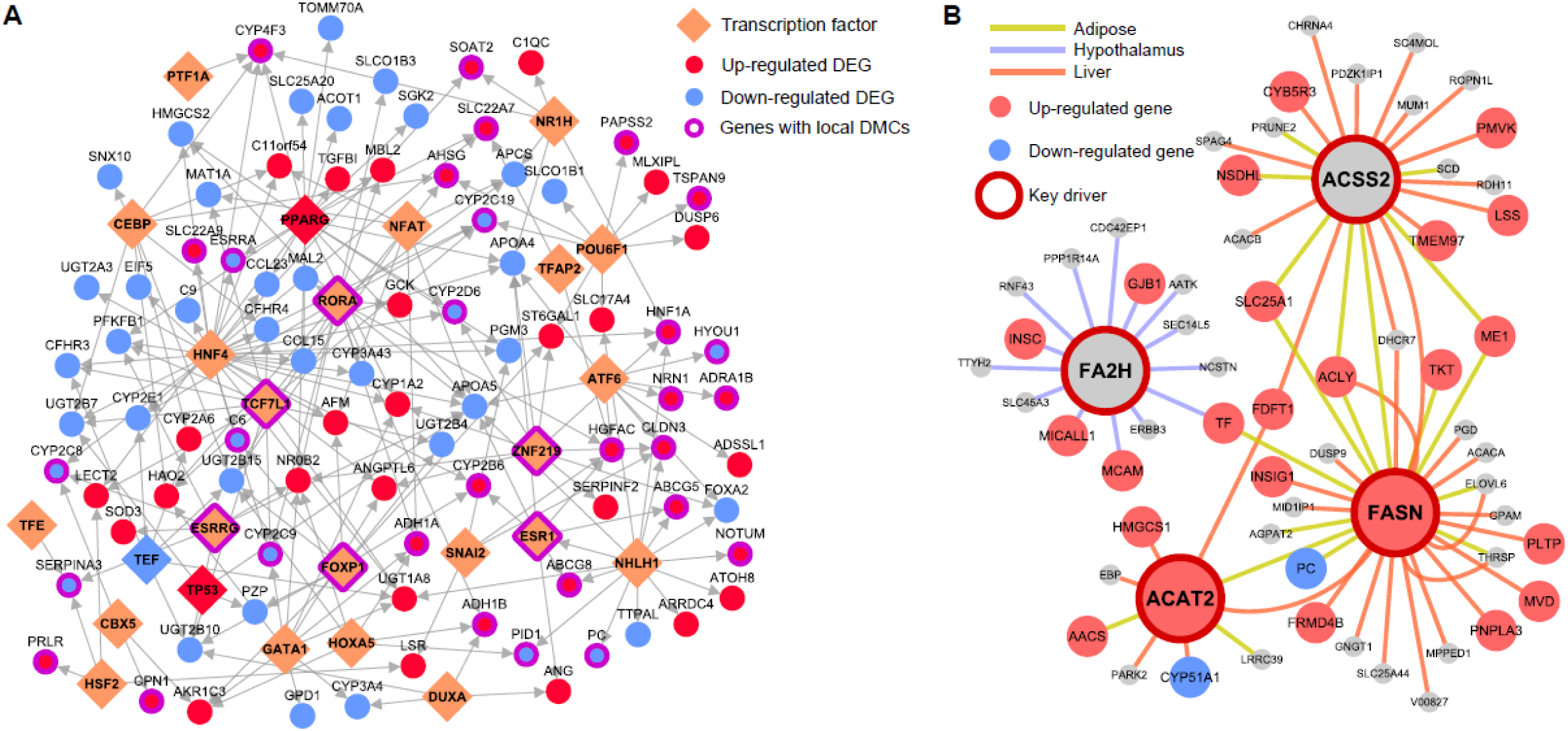
Transcription factors and key drivers orchestrate BPA induced gene expression level changes. (A) Liver transcription factor regulatory networks for the top ranked transcription factors (FDR < 5%) based on enrichment of liver DEGs among TF downstream targets. Network topology was based on FANTOM5. For TFs with > 20% overlapping downstream targets, only the TF with the lowest FDR is shown. (B) Gene-gene regulatory subnetworks (Bayesian networks) for cross-tissue key drivers. Network topology was based on Bayesian network modeling of each tissue using genetic and transcriptome datasets from mouse and human populations. For each tissue, if >= 2 datasets were available for a given tissue, a network for each dataset was constructed and a consensus network was derived by keeping only the high confidence network edges between genes (edges appearing in >= 2 studies).

### Identification of potential non-TF regulators governing BPA induced molecular perturbations

To further identify regulatory genes that mediate the action of BPA on downstream targets through non-TF mechanisms, we leveraged data-driven tissue-specific Bayesian networks (BNs) generated from multiple independent human and mouse studies (**S8 Table**). These data-driven networks are complementary to the TF networks used above and have proven valuable for accurately predicting gene-gene regulatory relationships and novel key drivers (KDs) [40-43, 61]. KDs were defined as network nodes whose surrounding subnetworks are significantly enriched for BPA exposure related DEGs. At FDR < 1%, we identified 21, 1, and 100 KDs in adipose, hypothalamus, and liver, respectively (**S9 Table**). The top KDs in adipose (top 5 KDs *Acss2, Pc, Agpat2, Slc25a1, Acly*), hypothalamus (*Fa2h*) and liver (top 5 KDs *Dhcr7, Aldh3a2, Fdft1, Mtmr11, Hmgcr*) were involved in cholesterol, fatty acid and glucose metabolism processes. In addition, three KDs, *Acss2* (Acetyl-Coenzyme A Synthetase 2), *Acat2* (Acetyl-CoA Acetyltransferase 2), and *Fasn* (Fatty Acid Synthase), were involved in the upregulation of DEGs in both adipose and liver, despite the fact that few DEG signatures overlap across tissues (**Fig 4B**). These KDs are consistent with the observed increased expression of several genes implicated in lipogenesis, including *Fasn*, and help explain the liver accumulation of triglycerides when mice are exposed to BPA [50]. Together, these results indicate that BPA may engage certain common regulators which have tissue-specific targets. The distinct upregulatory pattern within the subnetworks of individual KDs supports the potential functional importance of KDs in orchestrating the action of downstream genes. These KDs, along with the newly identified TFs from the above analysis, may represent novel regulatory targets which transmit the *in vivo* biological effects of BPA.

### BPA transcriptomic and methylomic signatures are related to metabolic traits in mice

To assess the relationship between the BPA molecular signatures and metabolic traits in the mouse model, the DEGs and DMCs from individual tissues were tested for correlation with the measured metabolic traits: body weight, free fatty acids (FFA), total cholesterol (TC), high density lipoprotein cholesterol (HDL), triglycerides (TG) and blood glucose. At p < 0.05, over two thirds of tissue-specific DEGs and over 60% DMCs were identified to be correlated with at least one metabolic trait (**Fig 5A, B**). Notably, liver DEGs exhibited stronger correlation with free fatty acid and triglycerides, whereas adipose DEGs were uniquely associated with glucose level, which is consistent with the pathway annotation results for these tissues. On the other hand, liver DMCs showed stronger correlations with metabolic traits than those from adipose and hypothalamus tissues.

**Fig 5.**
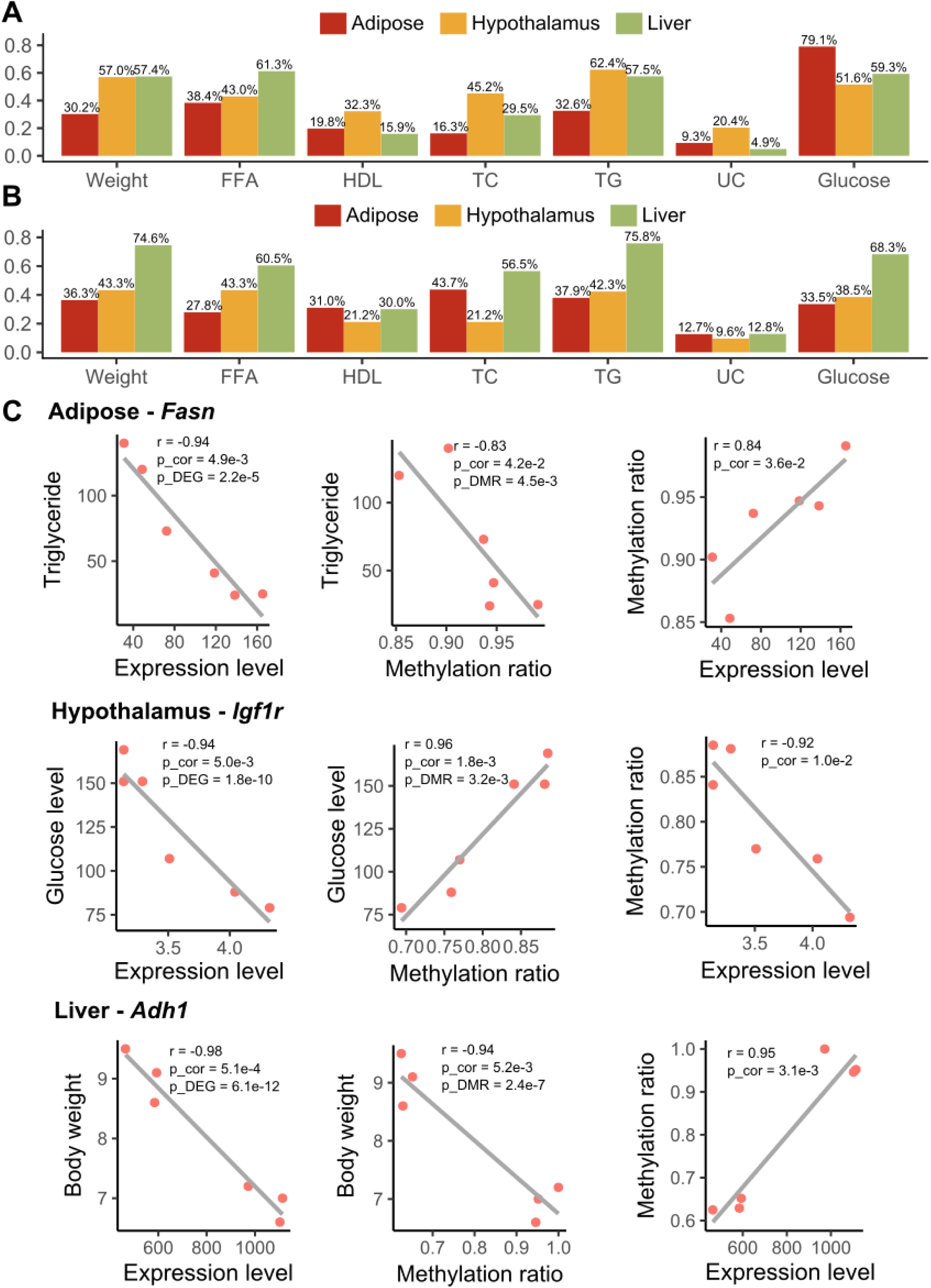
Correlation between gene expression, methylation and metabolic traits. (A) Percentage of tissue-specific DEGs that are correlated with metabolic traits (p < 0.05). (B) Percentage of tissue-specific DMCs that are correlated with metabolic traits (p < 0.05). (A-B) p-values were determined using Pearson correlation test. (C) Pair-wise correlation between expression level, methylation ratio and metabolic profiles (triglyceride, glucose level, body weight) for *Fasn, Igf1r and Adh1*. P_cor, p-value was determined using Pearson correction test; P_DEG was determined using differential expression test; P_DMC was determined using differential methylation test. Each dot represents a mouse.

Cross-examination of correlation across gene expression, DNA methylation, and metabolic traits revealed 35 consistent DEG-DMC-trait associations (3 in adipose, 4 in hypothalamus, and 28 in liver) (**S10 Table**). For example, in adipose tissue, *Fasn* (also a perturbed TF hotspot in adipose, and a shared KD in adipose and liver) was correlated with its exonic DMC at chr11:120816457, and both were correlated with triglyceride level; in hypothalamus, *Igf1r* (Insulin Like Growth Factor 1 Receptor) was correlated with its intronic DMC at chr7:68072768, and both were correlated with blood glucose level; in liver, *Adh1* (Alcohol Dehydrogenase 1A) was correlated with its intronic DMC at chr3:138287690, and both were correlated with body weight (**Fig 5C**). These results suggest that BPA alters local DMCs of certain genes to regulate gene expression, which may in turn regulate distinct metabolic traits.

### Relevance of BPA signature to human complex traits/diseases

Human observational studies have associated developmental BPA exposure with a wide variety of human diseases ranging from cardiometabolic diseases to neuropsychiatric disorders [14, 15, 62]. Large-scale human genome-wide association studies offer an unbiased view of the genetic architecture for various human traits/diseases, and intersections of the molecular footprints of BPA in our mouse study with human disease risk genes can help infer the potential disease-causing properties of BPA in humans. From the GWAS Catalog [63], we collected associated genes for 161 human traits/diseases (traits with fewer than 50 associated genes were excluded), and evaluated the enrichment for the trait associated genes among DEG and DMC signatures. At FDR < 5%, no trait was found to be significantly enriched for BPA DEGs. Surprisingly, despite the difference between tissue-specific DMCs (**Fig 3B**), 19 out the 161 traits showed consistently strong enrichment for DMCs across all three tissues at FDR < 1%. The top traits include body mass index (BMI) and type 2 diabetes (**Table 1**). As DNA methylation status is known to determine long-term gene expression pattern instead of immediate dynamic gene regulation, the BMI and diabetes associated genes may be under long-term programming by BPA-induced differential methylation, thereby affecting later disease risks.

**Table 1.** Top 5 human traits whose associated genes in genome-wide association studies are enriched for differentially methylated CpGs (DMCs) across adipose, hypothalamus and liver at FDR < 1% in MSEA.

| Human trait | Adipose |  | Hypothalamus |  | Liver |  |
| --- | --- | --- | --- | --- | --- | --- |
|  | P | FDR | P | FDR | P | FDR |
| Obesity-related traits | 1.28E-16 | 0.00% | 3.03E-15 | 0.00% | 2.71E-19 | 0.00% |
| Body mass index | 1.30E-13 | 0.00% | 3.74E-07 | 0.00% | 9.66E-12 | 0.00% |
| Post bronchodilator FEV1/FVC ratio | 8.17E-09 | 0.00% | 1.45E-08 | 0.00% | 3.67E-07 | 0.00% |
| Type 2 diabetes | 1.21E-05 | 0.03% | 8.97E-09 | 0.00% | 0.001243 | 0.92% |
| Platelet distribution width | 8.16E-08 | 0.00% | 7.62E-05 | 0.16% | 5.20E-05 | 0.12% |

The above analysis involving the GWAS catalog focused only on small sets of the top candidate genes for various diseases and may have limited statistical power. To improve the statistical power, we curated the full summary statistics from 61 human GWAS that are publicly available (covering millions of SNP-trait associations in each GWAS), which enabled us to extend the assessment of disease association by considering additional human disease genes with moderate to low effect sizes (**Methods**). This analysis showed that DEGs from all three tissues exhibited consistent enrichment for genes associated with lipid traits such as triglycerides, LDL, and HDL **(Fig 6A-C**). Interestingly, enrichment for birth weight and birth length was also observed for hypothalamus and liver signatures, respectively. Liver DEGs were also significantly associated with coronary artery disease, inflammatory bowel disease, Alzheimer’s disease, and schizophrenia. Top DEGs driving the inflammatory bowel disease association involve immune and inflammatory response genes (*PSMB9, TAP1, TNF*), whereas association with Alzheimer’s disease and schizophrenia involve genes related to cholesterol homeostasis (*APOA4, ABCG8, SOAT2*) and mitochondrial function (*GCDH, PDPR, SHMT2*), respectively. These results suggest that tissue-specific targets of BPA are connected to diverse human complex diseases through both the central nervous system and peripheral tissues.

**Fig 6.**
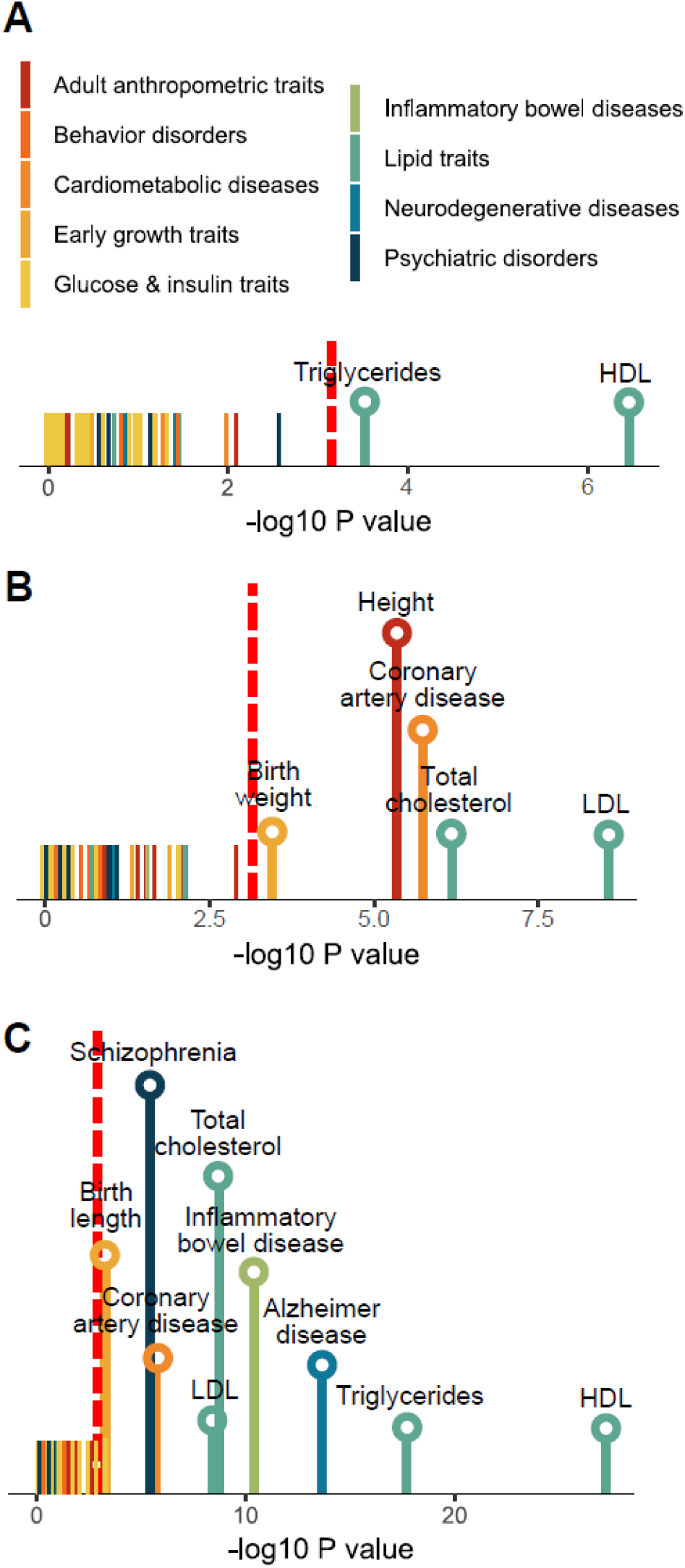
Association of differential expression signatures from adipose (A), hypothalamus (B) and liver (C) with 61 human traits/diseases, color coded into nine primary categories. P-values are determined using MSEA. Red dashed line indicates the cutoff for Bonferroni-corrected p = 0.05. Names of traits/diseases whose p-values didn’t pass Bonferroni-corrected cutoff were not shown.

## Discussion

This multi-tissue, multi-omics integrative study represents one of the first systems biology investigations of prenatal BPA exposure. By integrating systematic profiling of the transcriptome and methylome of multiple metabolic tissues with phenotypic trait measurements, large-scale human association datasets, and network analysis, we uncovered insights into the molecular regulatory mechanisms underlying the health effect of prenatal BPA exposure. Specifically, we identified tens to hundreds of tissue-specific DEGs and DMCs involved in diverse biological functions such as metabolic pathways (oxidative phosphorylation/TCA cycle, fatty acid, cholesterol, glucose metabolism, and PPAR signaling), extracellular matrix, focal adhesion, and inflammation (arachidonic acid), with DMCs partially explaining the regulation of DEGs. Network analysis helped reveal potential regulatory circuits post BPA exposure and pinpointed both tissue-specific and cross-tissue regulators of BPA activities, including TFs such as estrogen receptors, *PPARg*, and *HNF1A*, and non-TF key drivers such as *FASN*. Furthermore, the BPA gene signatures and the predicted regulators were found to be linked to a wide spectrum of disease-related traits in both mouse and human.

The large-scale disruption we observed in the transcriptome and methylome in adipose and liver was consistent with previous reports [33, 36, 64, 65]. Although our multi-tissue, multi-omics design limits the number of biological replicates we could have for each group, comparison of our results with previously published BPA studies with various study designs revealed consistent signals such as *Srebf1* in liver and a generally higher replication rate of the biological pathways between our study and the other studies than replication between the previously studies. Additionally, we focused our analyses on evaluating the aggregated behavior of BPA signatures using both pathway analysis and network modeling to reduce the potential noise and false positives at individual gene level, because the random chance to have multiple genes in the same pathway to be false positives is much lower. At the pathway level, we demonstrate the robustness of our results against published studies, supporting that relatively small sample size did not hinder our ability to uncover reliable biology despite variation at the individual signature level across studies. Moreover, our unique study design of examining multi-omics in multiple tissues in parallel yields higher comparability when integrating the results between data types and across tissues, as they were from the same set of animals and were profiled in the same conditions. Of note, the low consistency of DEGs across independent BPA studies points to the complexity of BPA actions under varying conditions, and warrants future large-scale coordinated investigations by the research community to systematically evaluate the molecular footprints of BPA.

Across all three tissues at the transcriptome level, we found that lipid metabolism and energy homeostasis related processes were consistently perturbed, with the scale of perturbation being strongest in liver. This aligns well with the significant changes in the plasma lipid profiles we observed in the offspring, the reported perturbation of lipid metabolism in fetal murine liver [65], and the reported susceptibility for nonalcoholic fatty liver diseases following BPA exposure [66-68]. The only shared gene across tissues, *Cyp51*, is involved in cholesterol and sterol biosynthesis and beta oxidation, and the only liver signature replicated across our and three previous studies and a top ranked TF regulator in our TF analysis, *Srebf1*, is a main regulator of lipid homeostasis, again supporting that metabolism is a central target of BPA. We also revealed an intriguing link between BPA and lncRNAs across tissues, whose functional importance in developmental processes, disease progression, and response to BPA exposure was increasingly recognized yet underexplored [69]. Our molecular data provides intriguing lncRNA candidates such as *Gm20319, Gm26917,* and *Yam1* for future in-depth functional analyses.

For adipose tissue, clusters of genes responsible for core histones were found to be uniquely altered. Along with the strong adipose-specific differential methylation status, our results revealed gonadal adipose tissues as an especially vulnerable site for BPA induced epigenetic reprogramming. Besides, developmental BPA exposure has been previously suggested to influence white adipocyte differentiation [70-72]. However, the adipocyte differentiation pathway was not significantly enriched in our study. This is consistent with the report by Angel et al. [72], where increased adipocyte number is only found in mouse offspring with prenatal BPA exposure at 5ug/kg/day and 500ug/kg/day, but not 5mg/kg/day. Additionally, we found significant enrichment for triglyceride biosynthesis and glucose metabolism genes at the differential methylation sites, suggesting that prenatal BPA exposure may affect fat storage and glucose homeostasis in the adipose tissue. Although here we mainly investigate gonadal adipose tissue as a surrogate for abdominal fat in the context of metabolic disorders, the information may be useful for exploring the relationship between this fat depot and the gonad.

With regards to the hypothalamus, our study is the first to investigate the effect of BPA on the hypothalamic transcriptome and DNA methylome. Hypothalamus is an essential brain region that regulates the endocrine system, peripheral metabolism, and numerous brain functions. We identified BPA-induced DEGs and DMCs that were enriched for extracellular matrix related processes such as axon guidance, focal adhesion, and various metabolic processes. These hypothalamic pathways have been previously associated with metabolic [41, 42] and neurodegenerative diseases [41, 73], and they could underlie the reported disruption of hypothalamic melanocortin circuitry after BPA exposure [74]. Our study highlights the hypothalamus as another critical yet under-recognized target for BPA.

By interrogating both the transcriptome and DNA methylome in matching tissues, we were able to directly assess both global and specific correlative relationships between DEGs and DMCs (**S5 Fig, Fig 3D**). Specifically, we found that DEGs are more likely to have correlated DMCs in the matching tissue, a trend that persists in non-promoter regions. Our results corroborate previous findings regarding the importance of gene body methylation in disease etiology [75, 76]. Given that over 90% of DMCs were found in non-promoter regions, closer investigation of the regulatory circuits involving these regions may unveil new insights into BPA response [52].

Known as an endocrine disrupting chemical, BPA has been speculated to exert its primary biological action by modifying the activity of hormone receptors, including estrogen receptors, PPARg and glucocorticoid receptors [55]. Indeed, the activity for the downstream targets of Pparg and three estrogen and estrogen-related receptors were found to be disrupted in liver by prenatal BPA exposure. More importantly, our unbiased data-driven analysis revealed many novel transcription factors and non-TF regulatory genes that also likely mediate BPA effects. In fact, many of the newly identified TF targets of BPA, such as Fasn, Srebf1, and several hepatic nuclear factors, showed much higher ranking in our regulator prediction analyses. In liver, a tightly inter-connected TF subnetwork was highly concentrated with BPA affected genes involved in metabolic processes such as cytochrome P450 system (*Cyp3a25, Cyp2a12, Cyp1a2*), lipid (*Apoa4, Abcg5, Soat2*) and glucose (*Hnf1a, Adra1b, Gck*) regulation, with extensive footprints of altered methylation status in the TFs and other subnetwork genes (**Fig 4A**). Therefore, our results support a widespread impact of BPA on liver transcriptional regulation, and the convergence of differential methylation and gene expression in this TF subnetwork implies that BPA perturbs this subnetwork via epigenetic regulation of the TFs, which in turn trigger transcriptomic alterations in downstream genes. In adipose, we discovered a regulatory axis governed by Nfya and Fasn that are known regulators of fatty acid metabolism and adipogenesis. NF-YA is a histone-fold domain protein that binds to the inverted CCAAT element in the *Fasn* promoter [60, 77], and both *Nfya* and *Fasn* were found to significantly perturbed by BPA in our study. Moreover, *Fasn* also serves as a cross-tissue KD, governing distinct groups of up-regulated lipid metabolism genes in adipose and liver post-BPA exposure (**Fig 4B**), supporting its role in mediating the BPA-induced lipid dysregulation at the systemic level. The significant correlation of gene expression and methylation for *Fasn* with triglyceride level furthers implicates its role as a network-level regulator and biomarker for BPA induced lipid dysregulation. Our observation of *Fasn* is consistent with evidences suggesting its susceptibility to methylation perturbation under obesogenic feeding [78] and its causal functional importance for fatty liver diseases [43, 79]. These novel regulators warrant future experimental testing of their causal regulatory role in BPA activities via genetic manipulation studies, such as knocking down or overexpressing *Fasn* to examine the modulation of BPA activities.

One unique aspect of this study is the linking of the molecular landscape of prenatal BPA exposure to traits/diseases in both mouse and human. In our mouse study, the observed changes in body weight, lipid profiles, and glucose level are highly concordant with the functions of the molecular targets. For instance, prenatal BPA exposure perturbs both the expression levels and the local DNA methylation status of *Fasn, Igf1r*, and *Adh1*. These DEGs and their local DMCs also significantly correlate with phenotypic outcomes, thus serving as examples of how DNA methylation and gene regulation bridge the gap between BPA exposure and phenotypic manifestation. To further enhance the translatability of our findings from mouse to human, we searched for human diseases linked to the BPA-affected genes. An intriguing discovery is the prominent overrepresentation of differential methylation signals in adipose, hypothalamus, and liver within known genes related to obesity and type 2 diabetes, supporting that BPA may impact obesity and diabetes risk through systemic reprogramming of DNA methylation. More sophisticated analysis incorporating the BPA differential gene expression and the full statistics of human genome-wide association studies corroborated the observed connection between prenatal BPA exposure and lipid homeostasis [80], birth weight [81], and coronary artery disease [14] reported in observational studies. Moreover, our findings suggest the involvement of prenatal BPA exposure in the development of inflammatory bowel syndrome, schizophrenia, and Alzheimer’s disease. These associations warrant future investigations.

One limitation of our work is the restriction of study scope to weaning age male mice with *in utero* BPA exposure below the NOAEL (5mg/kg/d) as a proof-of-concept for our systems biology framework. Considering that the effects of early-life exposure to BPA is highly variable and dependent on factors such as the dose, window, route, and frequency of exposure as well as genetic background, age, and sex [13], future studies testing these additional variables using large sample sizes are necessary to generate a comprehensive understandings of BPA risks under various exposure conditions.

## Conclusions

Our study represents the first multi-tissue, multi-omics integrative investigation of prenatal BPA exposure. The systems biology framework we applied revealed how BPA triggers cascades of regulatory circuits involving numerous transcription factors and non-TF regulators that coordinate diverse molecular processes within and across core metabolic tissues, thereby highlighting that BPA exerts its biological functions via much more diverse targets than previously thought. As such, our findings offer a comprehensive systems-level understanding of tissue sensitivity and molecular perturbations elicited by prenatal BPA exposure, and offer promising novel candidates for targeted mechanistic investigation as well as much-needed network-level biomarkers of prior BPA exposure. The strong influence of BPA on metabolic pathways and cardiometabolic phenotypes merits it characterization as a general metabolic disruptor posing systemic health risks.

## Methods

### Ethics statement

All animal experiments were performed in accordance with the Institutional Animal Care and Use Committee (IACUC) guidelines. Animal studies and procedures were approved by the Chancellor’s Animal Research Committee of the University of California, Los Angeles (Protocol #2012-059-21).

### Mouse model of prenatal BPA exposure

Inbred C57BL/6 mice were maintained on a special diet 5V01 (LabDiet), certified to contain less than 150ppm estrogenic isoflavones, and housed under standard housing conditions (room temperature 22–24°C) with 12:12 hr light:dark cycle before mating at 8-10 weeks of age. Upon mating, female mice were randomly assigned to either the BPA treatment group or the control group. From 1-day post-conception (dpc) to 20 dpc, BPA (Sigma-Aldrich, St. Louis, MO) dissolved in corn oil was administered to pregnant female mice via oral gavage (mimicking common exposure route in humans) at 5mg/kg/day on a daily basis. Control mice were fed the same amount of empty vehicle. BPA exposure was restricted to experimental manipulation through the use of polycarbonate-free water bottles and cages. Offspring from each treatment were maintained on a standard chow diet (Newco Distributors Inc, Rancho Cucamonga, CA). Offspring in the vehicle- and BPA-treated groups were derived from 3 and 4 litters by different dams, respectively, to help assess and adjust for litter effects.

### Characterization of cardiometabolic phenotypes and tissue collection

Body weight of offspring was measured daily from postnatal day 5 up to the weaning age of 3 weeks. Mice were fasted overnight before sacrifice, and plasma samples were collected through retro-orbital bleeding. Serum lipid and glucose traits including total cholesterol, HDL, un-esterified cholesterol (UC), TG, FFA, and glucose were measured by enzymatic colorimetric assays at UCLA GTM Mouse Transfer Core as previously described [41]. Gonadal white adipose tissue, hypothalamus, and liver tissues were collected from each animal, flash frozen in liquid nitrogen, and stored at –80°C. For white adipose tissue, we chose the gonadal depot mainly due to its similarity to abdominal fat, established relevance to cardiometabolic risks, tissue abundance, and the fact that it is the most well-studied adipose tissue in mouse models. All mouse experiments were conducted in accordance with and approved by the Institutional Animal Care and Use Committee at University of California, Los Angeles.

### RNA sequencing (RNA-seq) and data analysis

A total of 18 RNA samples were isolated from gonadal adipose, hypothalamus and liver tissues (n = 3 per group per tissue; for each group, mice were randomly selected from litters of different dams in independent cages) from male offspring using the AllPrep DNA/RNA Mini Kit (QIAGEN GmbH, Hilden, Germany). We focused on profiling male tissues because of stronger phenotypes observed in males (**Fig 1B-E**). Samples were processed for library preparation using TruSeq RNA Library Preparation Kit (Illumina, San Diego, CA) for poly-A selection, fragmentation, and reverse transcription using random hexamer-primers to generate first-strand cDNA. Second-strand cDNA was generated using RNase H and DNA polymerases, and sequencing adapters were ligated using the Illumina Paired-End sample prep kit. Library products of 250-400bp fragments were isolated, amplified, and sequenced with Illumina Hiseq2500 System. After quality control using FastQC [82], the HISAT-StringTie pipeline [83] was used for sequence alignment and transcript assembly. Identification of DEGs were conducted using DEseq2 [84]. To account for multiple testing, we used the q-value method [85]. After excluding genes with extremely low expression levels (FPKM < 1), only DEGs demonstrating differential expression comparing the BPA and control groups per tissue at an FDR < 5% were used for biological pathway analysis, network analysis, and phenotypic data integration, as described below.

### Reduced representation bisulfite sequencing (RRBS) and data analysis

We constructed RRBS libraries for 18 DNA samples from adipose, hypothalamus and liver tissues from male offspring (n = 3 per group per tissue from the same set of tissues chosen for transcriptome analysis described above). The DNA samples were quantified using the dsDNA BR assay (Qubit, Waltham, MA) and 100ng of DNA was used for library preparation. After digestion of the DNA with the MspI enzyme, samples underwent an end-repair and adenylation process, followed by adapter ligation using the Truseq barcode adapter (Illumina, San Diego, CA), size selection using AMPure Beads (Beckman Coulter, Brea, CA), and bisulfite treatment using the Epitect Kit (Qiagen, Germantown, MD). Bisulfite-treated DNA was then amplified using the Truseq Library Prep Kit (Illumina, San Diego, CA) and sequenced with the Illumina Hiseq2500 System. Bisulfite-converted reads were processed and aligned to the reference mouse genome (GRCm38/mm10 build) using the bisulfite aligner BSMAP [86]. We then used MOAB [87] for methylation ratio calling and identification of DMCs. FDR was estimated using the q-value approach. Loci with methylation level changes of > 5% between BPA and control groups and FDR < 0.05 for each tissue were considered statistically significant DMCs. To annotate the locations of the identified DMCs in relation to gene regions and repetitive DNA elements accessed from UCSC genome browser, we used the Bioconductor package “annotatr” [88]. Specifically, gene regions were categorized into 1) 1-5kb upstream of the transcription start site (TSS), 2) promoter (< 1kb upstream of the TSS), 3) 5’ untranslated region (UTR), 4) exons, 5) introns, and 6) 3’UTR. The “annotatr” package was also used to annotate DMCs for known lncRNAs based on GENCODE Release M16. Over-representation of DMCs within each category was calculated using Fisher’s exact test. We further evaluated the link between DEGs and their local DMCs (DMCs annotated as any of the 6 above mentioned gene regions) by correlating the methylation ratio of DMCs with the expression level of DEGs.

### Pathway, network, and disease association analyses of DEGs and DMCs using the Mergeomics R package

To investigate the functional connections among the BPA-associated DEGs or DMCs (collectively referred to as molecular signatures of BPAs) and to assess the potential association of BPA affected genes with diseases in human populations, we utilized the Mergeomics package [49], an open-source bioconductor package (https://bioconductor.org/packages/devel/bioc/html/Mergeomics.html) designed to perform various integrative analyses in multi-omics studies. Mergeomics consists of two main libraries, Marker Set Enrichment Analysis (MSEA) and Weighted Key Driver Analysis (wKDA). In the current study, we used MSEA to assess 1) whether known biological processes, pathways or transcription factor targets were enriched for BPA molecular signatures as a means to annotate the potential functions or regulators of the molecular signatures, and 2) whether the BPA signatures demonstrate enrichment for disease associations identified in human genome-wide association studies (GWAS) of various complex diseases (**S7 Fig**). wKDA leverages gene network topology (interactions or regulatory relations among genes) and edge weight (strength or reliability of interactions and regulatory relations) information of graphical gene networks to predict potential key regulators of a given group of genes, in this case, the BPA-associated DEGs (**S8 Fig**). Both MSEA and wKDA were built around a chi-square like statistics (**S1 Text**) that yields robust findings that have been experimentally validated [42, 43, 49]. Details of each usage of the Mergeomics package are discussed below.

### Functional annotation of DEGs and DMCs

To infer the functions of the DEGs and DMCs affected by BPA, we used MSEA to annotate the DEGs or local genes adjacent to the DMCs with known biological pathways curated from the Kyoto Encyclopedia of Genes and Genomes (KEGG) [89] and Reactome [90]. In brief, we extracted the differential expression p-values of genes in each pathway from the differential expression or methylation analyses and compared these p-values against the null distribution of p-values from random gene sets with matching gene numbers. If genes in a given pathway collectively show more significant differential expression or differential methylation p-values compared to random genes based on a chi-square like statistic, we annotate the DEGs or DMCs using that pathway (**S1 Text**). DEGs and DMCs can have multiple over-represented pathways.

### Identification of transcription factor hotspots perturbed by BPA

To dissect the regulatory cascades of BPA, we first assessed whether BPA-associated DEGs were downstream targets of specific transcription factors. The hypothesis behind this analysis is that BPA first affects TFs which in turn regulate the expression of downstream genes. We used TF regulatory networks for adipose, brain, and liver tissue retrieved from the FANTOM5 database [59]. Note that only a whole brain (instead of hypothalamus) TF network was available, which may only partially represent hypothalamic gene regulation. Each TF network was processed to keep the edges with high confidence (**S1 Text**). To identify TFs whose targets were perturbed by BPA, the downstream nodes of each TF in the network were pooled as the target genes for that TF. We then assessed the enrichment for BPA exposure related DEGs among the target genes of each TF using MSEA. TFs with FDR < 5% were considered statistically significant. Cytoscape software was used for TF network visualization [91].

### Bayesian network and Weighted Key Driver Analysis (wKDA) to identify potential non-TF regulators

To further identify non-TF regulators that sense BPA and then perturb downstream genes, we used Bayesian networks (BN) of adipose, hypothalamus and liver tissues constructed from genetic and transcriptomic data from several large-scale mouse and human studies (**S1 Text and S8 Table**). wKDA was used to identify network key drivers (KDs), which are defined as network nodes whose neighboring subnetworks are significantly enriched for BPA-associated DEGs. Briefly, wKDA takes gene set G (i.e. BPA DEGs) and directional gene network N (i.e. BNs) as inputs. For every gene K in network N, neighboring genes within 1-edge distance were tested for enrichment of genes in G using a chi-square like statistics followed by FDR assessment by permutation (**S1 Text and S8 Fig**). Network genes that reached FDR < 0.05 were reported as potential KDs.

### Association of BPA DEGs and DMCs with mouse phenotypes and human diseases/traits

To assess whether the BPA molecular signatures were related to phenotypes examined in the mouse offspring, we calculated the Pearson correlation coefficient among expression level of DEGs, methylation ratio of DMCs, and the measurement of metabolic traits. For human diseases or traits, we accessed the GWAS catalog database [63] and collected the lists of candidate genes reported to be associated with 161 human traits/diseases (P < 1e-5). These genes were tested for enrichment of the BPA DEGs and DMCs in our mouse study using MSEA. We further curated all publicly available full summary statistics for 61 human traits/diseases from various public repositories (**S1 Text and S11 Table**). This allowed us to apply MSEA to comprehensively assess the enrichment for human disease association among BPA transcriptomic signatures using the full-spectrum of large-scale human GWAS. For each tissue-specific gene signature, we used the SNPs within a 50kb chromosomal distance as the representing SNPs for that gene. The trait/disease association p-values of the SNPs were then extracted from each GWAS and compared to the p-values of SNPs of random sets of genes to assess whether the BPA signatures were more likely to show stronger disease association in human GWAS (**S1 Text and S7 Fig**). This strategy has been successfully used in our previous animal model studies to assess the connection of genes affected by environmental perturbations such as diets and trauma to various human diseases [41, 92].

## Acknowledgments

LS is supported by UCLA Dissertation Year Fellowship, Eureka Scholarship, Hyde Scholarship, Burroughs Wellcome Fund Inter-School Program in Metabolic Diseases Fellowship, and China Scholarship Council. XY is supported by NIH DK104363 and NS103088, and Leducq Foundation. We would also like to thank Zhe Ying’s assistance in collecting mice hypothalamus tissue, and Dr. Guanglin Zhang’s assistance in the RRBS experiments.

## Competing interests

The authors declare that they have no competing interest.

## Supporting information captions

S1 Text. Supplemental Methods.

S1 Fig. Body weight of male and female offspring mice at weaning age by litters. Red arrows indicate offspring male mice selected for molecular profiling.

S2 Fig. Prenatal BPA exposure induced expression change for genes from the adipocyte differentiation, triglyceride biosynthesis, glucose metabolism, and core histone genes in the adipose tissue. P-values for enrichment of pathway genes among DEGs (shown in parenthesis in each panel heading) were determined by MSEA. *p < 0.05 in differential expression tests for individual genes by DEseq2; **FDR < 5% in differential expression tests for individual genes by DEseq2.

S3 Fig. Comparison of the liver DEGs and their functional annotations against published datasets in GEO. (A) Descriptions of the study design of different datasets. (B) Venn Diagram of the DEGs identified in different datasets. DEGs were determined by Limma at p < 0.01. (C) The percentage of DEGs (from the datasets in each row header) that are replicated (by the datasets in each column header). Numbers in parenthesis indicate the percentage of DEGs that are replicated by at least one independent study, and the significance of the replication percentage determined by permutation test. (D) Venn Diagram of the functional annotations for the DEGs identified in different datasets. Functional annotations were determined by MSEA at FDR < 5%. (E) The percentage of functional annotations (from the datasets in each row header) that are replicated (by the datasets in each column header). Numbers in parenthesis indicate the percentage of annotations that are replicated by at least one independent study, and the significance of the replication percentage determined by permutation test.

S4 Fig. Gene body location distribution for hyper- and hypo-methylated DMC s in adipose, hypothalamus, and liver.

S5 Fig. Quantile-quantile plots for the absolute Pearson correlation with local DMC for DEGs and Non DEGs in adipose, hypothalamus, and liver tissue. Statistical difference of the distribution of correlation value between DEGs (FDR < 5%) and non DEGs is determined by the Kolmogorov–Smirnov test.

S6 Fig. Scatter plots of correlations between DEG expression levels and DMC methylation ratios for *Slc25a1* in adipose, *Mvk* in hypothalamus, and *Gm20319* in liver.

S7 Fig. Schematic illustration of MSEA

S8 Fig. Schematic illustration (A) and key driver identification algorithms (B) of wKDA.

S1 Table. List of DEGs with p < 0.05 in adipose, hypothalamus and liver tissue following prenatal exposure to BPA.

S2 Table. Count of DEGs in adipose, hypothalamus and liver tissue following prenatal exposure to BPA.

S3 Table. Results of functional annotation of DEGs in in adipose, hypothalamus and liver tissue following prenatal exposure to BPA

S4 Table. Count of differentially methylated regions in hypothalamus and liver tissue following prenatal exposure to BPA.

S5 Table. Results of functional annotation of DMCs in in adipose, hypothalamus and liver tissue following prenatal exposure to BPA.

S6 Table. List of pairs of DEGs and local DMCs with significant correlation (p < 0.05) in expression level and methylation ratio

S7 Table. List of transcription factors whose downstream targets were significantly enriched for DEGs (p < 0.05).

S8 Table. Data resources and references for the construction of Bayesian gene-gene regulatory networks.

S9 Table. List of tissue-specific key drivers with FDR < 1%

S10 Table. List of DEGs with significant correlation between expression level, methylation ratio of local DMCs, and cardiometabolic traits (p < 0.05).

S11 Table. Source of publicly available full summary-level statistics from human genome-wide association studies.

